# MR-PET head motion correction based on co-registration of multi-contrast MR images

**DOI:** 10.1101/450676

**Authors:** Zhaolin Chen, Francesco Sforazzini, Jakub Baran, Thomas Close, N. Jon Shah, Gary F. Egan

**Affiliations:** Monash Biomedical Imaging, Monash University, Australia; Department of Electrical and Computer Systems Engineering, Monash University, Australia; Department of Biophysics, Faculty of Mathematics and Natural Sciences, University of Rzesow, Poland; Australian National Imaging Facility, Australia; Institute of Neuroscience and Medicine – 4, Forschungszentrum Jülich GmbH, Jülich, Germany; Australian Research Council Centre of Excellence for Integrative Brain Function, Monash University, Australia; Monash Institute of Cognitive and Clinical Neuroscience, Monash University, Australia

**Keywords:** Simultaneous MR-PET, PET/MR, PET motion artefacts, PET motion correction, MR image registration, multiple acquisition frame (MAF), MR-guided motion correction, MR-guided MAF

## Abstract

Head motion is a major source of image artefacts in neuroimaging studies and can lead to degradation of the quantitative accuracy of reconstructed PET images. Simultaneous Magnetic Resonance-Positron Emission Tomography (MR-PET) makes it possible to estimate head motion information from high-resolution MR images and then correct motion artefacts in PET images. In this paper, we introduce a fully automated PET motion correction method, MR-guided MAF, based on the co-registration of multi-contrast MR images. The performance of the MR-guided MAF method was evaluated using MR-PET data acquired from a cohort of ten healthy participants who received a slow infusion of fluorodeoxyglucose ([18-F]FDG). Compared with conventional methods, MR guided PET image reconstruction can reduce head motion introduced artefacts and improve the image sharpness and quantitative accuracy of PET images acquired using simultaneous MR-PET scanners. The fully automated motion estimation method has been implemented as a publicly available web-service.

## Introduction

The lengthy duration of the simultaneous Magnetic Resonance-Positron Emission Tomography (MR-PET) brain imaging experiments can lead to head motion induced artefacts in the PET images (Chen, et al., 2018). Even sub-millimetre motion which is not manifest as visible image artefacts can result in systematic and regionally specific biases in MRI anatomical estimations (Alexander-Bloch et al., 2016). Significant effects are observed in fMRI functional connectivity measurements due to head motion (Satterthwaite et al., 2012). With recent improvements in PET scanner resolution, head motion is increasingly becoming one of the major causes of image quality degradation, including reduction of spatial resolution and erroneous estimation of radio-ligand concentrations.

A widely used technique for correcting head motion in PET and PET-CT scanners is the multiple acquisition frame (MAF) method (Picard and Thompson, 1997). The MAF method subdivides the PET raw data (i.e. list-mode data) into a number of short duration temporal frames, with the frames then co-registered to correct for head motion under the assumption that intra-frame motion is negligible.

External motion tracking devices can further be installed to monitor motion (see (Maclaren et al., 2013) for a detailed review). These devices can provide excellent motion estimation accuracy and temporal sampling of motion parameters at milliseconds temporal resolution, but in general they are complex to setup and often have patient compliance issues. Additionally, MR compatibility and PET attenuation aspects of an external device have to be considered thoroughly for application to a hybrid MR-PET scanner. Because of the complexity in workflow and potential patient compliance issue, external device motion correction methods are currently not commonly used in routine clinical and experimental studies.

Data-driven methods form another category of motion correction methods in PET imaging. Recently, Thielemans et al. applied the principal component analysis (PCA) method to detect head movements directly from PET sinogram or list-mode data, and then used the estimated motion position information to guide the MAF framing process (Schleyer et al., 2015; Thielemans et al., 2013). The PCA motion detection method is based on the identification of changes in the principal components of the PET time activity curve, and compared with the conventional MAF technique, the PCA guided MAF has demonstrated to reduce intra-frame motion in PET image reconstruction. However, the PCA motion detection can only work when the tissue biological kinetics are stable and therefore any signal change in the tissue time activity curve is due to motion. This assumption is invalid in dynamic PET data acquisition where tracer uptake in the brain increases with time. Furthermore, PET based methods rely on co-registration of PET images which have intrinsic lower spatial resolution and anatomical contrast compared with MR images.

Recently, slow infusion based dynamic fluorodeoxyglucose ([18-F]FDG) PET imaging has shown promising results for investigating dynamic brain metabolism (Hahn et al., 2016; Villien et al., 2014). In these methods, PET list-mode data are acquired for 60-90 minutes and then binned into 1-min frames. Due to the long acquisition time, motion correction is critical in the dynamic PET imaging. Conventional PET data-driven approaches cannot accurately estimate the motion since the radioactivity distribution in the brain accumulates and changes over time. Therefore, a reliable motion correction method is still required.

Simultaneous MR-PET makes it possible to model head motion from high-resolution MR images and then correct motion artefacts in PET images. While PET data driven methods may work for [18-F]FDG PET when signal to noise ratio (SNR) is sufficient, the MR based motion correction can be advantageous in many applications including low dose FDG PET and other tracers such as receptor-targeted PET where spatial SNR is limited. Echo Planar Imaging (EPI) MRI volumes are often used as image navigators to track head movements. In Blood-Oxygen-Level Dependent functional MRI (BOLD fMRI) experiments, dynamic EPIs are acquired in every repetition time (TR=2 seconds or less), which provide motion estimates for PET data correction (Catana et al., 2011; Ullisch et al., 2012). Single EPI volumes can also be inserted in between MR sequences/scans which are often several minutes apart (Keller et al., 2015). The specific advantages of using EPI navigators to perform motion correction include their high temporal resolution (i.e. in seconds) and spatial resolution and SNR for accurate image registration. Many software toolboxes (e.g. FSL, SPM, ANTS, and etc) have been introduced to co-register fMRI EPI volumes, and these software tools have achieved excellent image co-registration accuracy. Ardekani et al., (2001) demonstrated that excellent image co-registration accuracy in the order of 0.20 mm can be obtained when the image SNR is greater than 5. However, inserting EPI acquisitions amongst MR sequences takes additional imaging time. In recent work, EPI navigators have also been embedded directly into T1 weighted MR sequences (i.e. in every repetition time TR) for intra-sequence motion correction (Tisdall et al., 2016). However, this method adds additional acquisition time to the minimum TR and is not routinely available for other MR sequences.

Our aim in this research was to develop a fully automated MR guided method based on co-registration of multi-contrast MR images with different resolution and imaging parameters. The MR guided PET motion correction method has the following advantages: 1) a fully automated method that does not require any image or k-space navigators, 2) provides excellent motion estimation accuracy due to the high spatial resolution of the MR images, and 3) it can be applied in both static and dynamic PET experiments. The introduced method, MR-guided MAF, optimises the MR image registration for all types of MR image contrasts (e.g. T1, T2, EPI BOLD, Diffusion Weighted Imaging (DWI), Arterial Spin Labelling (ASL), and etc). The inclusion of different MR image contrasts makes it possible to correct motion during the complete neuroimaging examination. Similar to BOLD EPIs, DWI and ASL sequences are also dynamic scans and can be used to extract motion estimates with high temporal resolution. Anatomical T1 and T2 weighted MR acquisitions normally have long scanning duration (e.g. 5-10 mins), and motion estimates from them can be limited in temporal resolution. Nevertheless, in a comparable approach Keller et al (2015) demonstrated improved PET image quality using navigators which are several minutes apart. The MR-guided MAF extracts motion parameters which are then used to rebin PET raw data into a multiple acquisition frame reconstruction. The PET attenuation map can also be re-aligned to match the head position for each frame to further improve the quality of the PET image reconstruction. The MR-guided MAF method was evaluated on a volunteer with controlled head motion as well as on a cohort of ten subjects who were instructed to minimise their head motion during data acquisition. Participants were slowly administered 260MBq [18-F]FDG PET at constant infusion rate, and to the best of our knowledge, the impact of MR based PET motion correction has not been previously investigated in a cohort of subjects undergoing a slow infusion FDG PET experiment. The motion correction performance of the MR-guided MAF method was evaluated using a static (single frame) PET image reconstruction in the cohort. The impact of the PET attenuation map re-alignment was investigated in the motion controlled experiment. Improvements in the image sharpness and accuracy of PET image quantification during the dynamic PET image reconstruction were also investigated.

## Methods

The MR-guided MAF method contains two main steps: i) multi-contrast MR co-registration and ii) the MR-guided MAF PET image reconstruction. In the first step, the motion parameters are estimated by image registration and concatenation of transformation matrices. These motion parameters are then used to guide the reconstruction of static or dynamic PET images in the second step. An overview of the MR-guided MAF method is shown in Figure 1.

**Figure 1.**
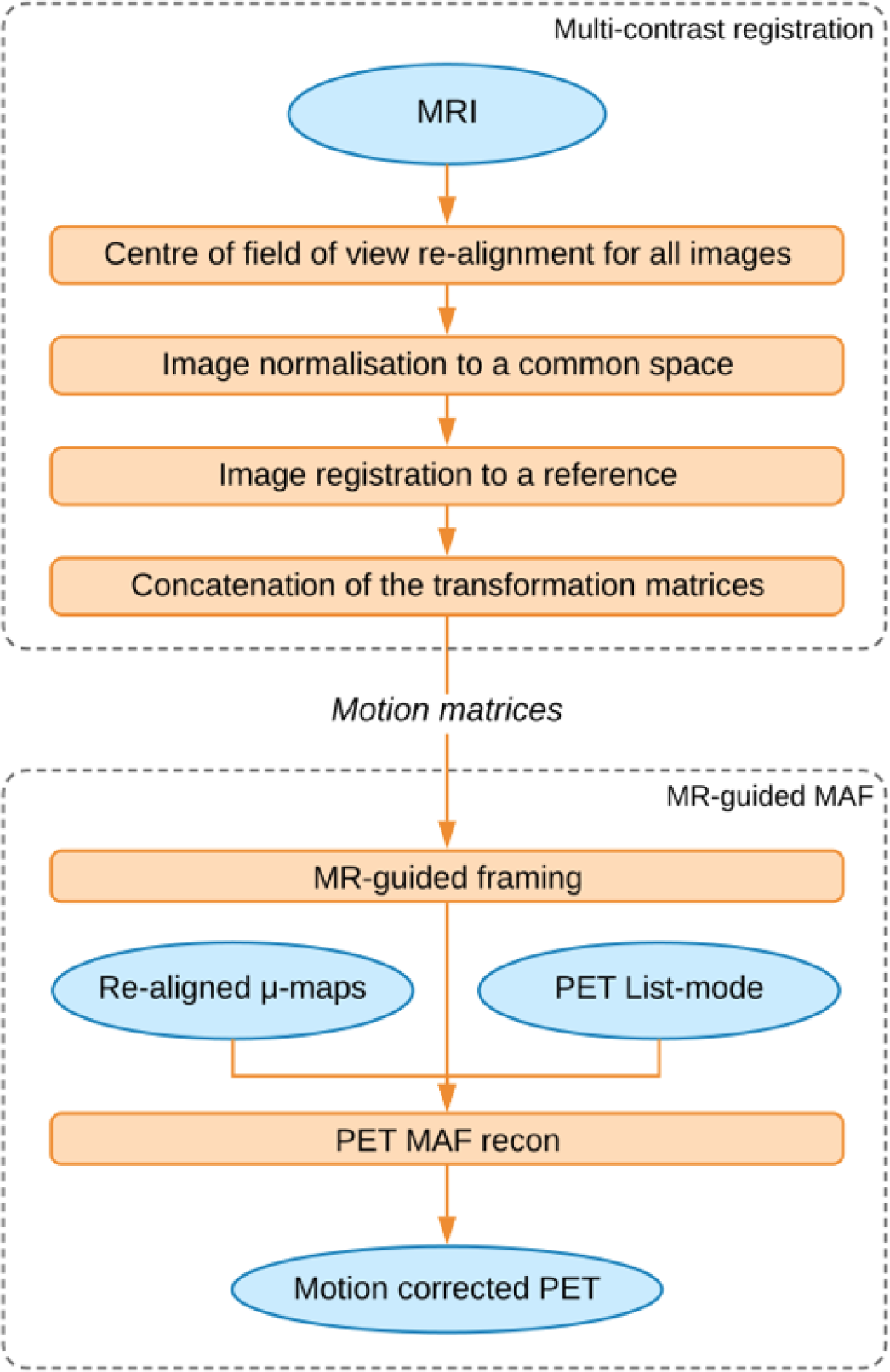
Overview of the MR-guided MAF method.

### Motion estimation based on multi-contrast MR image registration

#### Selection of reference image for registration

A reference image was first selected for registration of the multi-contrast MR images. The selected reference image was chosen to minimize the bias introduced by image registration. T1-weighted contrasts are typically used as the anatomical reference due to their good grey/white matter contrast. In this work, we compared the registration imprecision from both 3D isotropic T1-weighted and T2-weighted images as references.

In order to determine the optimal anatomical reference image between the T1 and T2 weighted images, an IIDA brain phantom (Iida et al., 2013) was used to acquire T1 weighted 3D MPRAGE (Magnetization-Prepared Rapid Gradient-Echo), T2 weighted 3D FLAIR (Fluid-Attenuated Inversion Recovery), PD (Proton Density weighted), DWI (Diffusion Weighted Imaging) based on Spin Echo-Echo Planar Imaging (SE-EPI) and BOLD fMRI (Blood-Oxygen-Level Dependent functional MRI) using Gradient Echo EPI (GE-EPI) and 2D GRE (Gradient Echo) (see Table 1 for detailed acquisition parameters). The brain phantom was fixed inside the head coil and kept free of motion during the acquisition. Each MR image contrast was then rigidly (6 degree of freedom) registered to the T1 and T2 weighted images. The acquisition was repeated four times to calculate the standard errors of the mean registration bias. In this work, we selected T2 weighted FLAIR image as the reference image (see Results section).

**Table 1:** MR image acquisition parameters for the motion instructed volunteer and the phantom experiments.

| **Scan** | **TR** | **TE** | **matrix** | **slices** | **resolution** |
| --- | --- | --- | --- | --- | --- |
| **UTE** | 11.94 | 0.07, 2.46 | 192x192 | 192 | 1.5x1.5x1.5 |
| **T1 MPRAGE** | 1640 | 2.34 | 256x256 | 176 | 1.0x1.0x1.0 |
| **T2 FLAIR** | 3200 | 418 | 256x256 | 176 | 1.0x1.0x1.0 |
| **Proton density** | 3800 | 16 | 128x128 | 34 | 1.0x1.0x2.0 |
| **ASL** | 2500 | 13 | 64x64 | 9 | 4.0x4.0x10.0 |
| **BOLD fMRI** | 3200 | 20 | 64x64 | 60 | 3.5x3.5x3.0 |
| **DWI** | 13100 | 110 | 96x96 | 60 | 2.5x2.5x2.5 |
| **2D GRE** | 466 | 4.92, 7.38 | 64x64 | 44 | 3.0x3.0x3.0 |

TR: Repetition Time [ms]; TE Echo Time [ms]; Res: Resolution [mm<sup>3</sup>].

#### Pre-processing of images

To improve image registration accuracy and robustness, brain extraction (BET, Smith 2002) was applied to each MR image.

For EPI acquisitions (i.e. BOLD and diffusion weighted), a geometric distortion correction was applied using an opposite-phase encoding EPI image. The distortion correction was implemented using the FSL-TOPUP toolkit (Andersson et al., 2003).

The T2 weighted image was segmented into grey and white matters and CSF using FAST (Zhang et al., 2001). The segmented white matter boundaries were used to improve the registration accuracy of the BOLD and ASL (Arterial Spin Labelling) MR images to the reference T2 weighted images.

#### Multi-contrast image registration

Motion matrices containing the rotational and translational parameters for the anatomical MR images (i.e. T1, T2, and PD) and multi-volume images (i.e. BOLD, ASL and DWI) were estimated as per the following steps:

1) All MR images (the first volume if a multi-volume MRI) were normalized to the image space of the reference, accounting for field-of-view and images resolution differences.
2) a) For anatomical MRI, each image contrast was rigidly registered to the T2 weighted reference image using FSL FLIRT (Jenkinson and Smith, 2001).
b) For multi-volume MRI, only the first image volume was rigidly registered to the reference. The registration steps for BOLD fMRI and ASL were optimised using white matter boundaries from the T2 weighted images. Each of the remaining volumes were then aligned to the first volume using MCFLIRT (Jenkinson et al., 2002) for ASL and BOLD fMRI, and using EDDY (Andersson and Sotiropoulos, 2016) for DWI (with a b0 volume as the first volume).
3) a) For anatomical MRI, the motion matrix of the corresponding image contrast was calculated by multiplying the inverse of the transformation matrices in step 1) and the registration matrices in step 2a).
b) For multi-volume MRI, the motion matrix of the corresponding image volume was calculated by multiplying the inverse of each of the registration matrix obtained in step 2b).

Rotational and translational parameters, as well as the mean displacement, were derived from the estimated motion matrices. During MR idling times, the last known motion estimates were used. For the anatomical MRI images (e.g. T1 and T2 weighted) which take several minutes to acquire, the estimated motion parameters represent an estimate of the averaged motion throughout the acquisition period.

### Multiple Acquisition Frame (MAF) correction

#### MR guided MAF

The mean displacement parameter was used to guide the MAF algorithm in the subdivision of the PET list-mode data into multiple motion correction frames. Specifically, the following two criteria were used to form a new motion correction frame when motion occurred:

- The absolute difference between the mean displacement values of two consecutive volumes was greater than a predefined threshold parameter (*d_1_*). This criterion determined whether a sudden movement occurred. The parameter *d_1_* was set to 2 mm in this work.
- Mean displacement between the current volume and the reference was greater than a predefined threshold parameter (*d_2_*). This criterion determined whether a gradual motion (e.g. the subject’s head position is slowly drifting) occurred. The parameter *d_2_* was also set to 2 mm in this work.

The choices of the threshold *d_1_* and *d_2_* values were a compromise between the computational time and the motion estimation accuracy, with 2mm chosen because it was less than the voxel size of the reconstructed PET images. Furthermore, the minimum duration for a motion correction frame was one minute to ensure the motion correction frame had sufficient radioactivity counts to reconstruct a PET image.

<u>μ*-map realignment:*</u> The attenuation correction μ-map was re-aligned to the head position for each motion correction frame prior to the PET image reconstruction. All the MR derived motion parameters within one motion correction frame were averaged to obtain an averaged motion estimate which was then applied to the original μ-map data.

Timestamps, re-aligned attenuation maps and PET list-mode data were used with the PET image reconstruction software (see section 2.3) to reconstruct one PET image per motion correction frame. The final motion corrected static PET image was calculated using the frame duration weighted average of the motion corrected PET images. The dynamic PET images were reconstructed by re-binning the list-mode data into 90 frames (one minute per frame), and the averaged motion parameters within each of these frames were used to correct for head motion.

#### Fixed frame MAF

The fixed frame MAF is often used to compare motion correction methods (Schleyer et al., 2015). In the fixed frame MAF, the PET list-mode data were first re-binned into one minute length frames, which were then reconstructed and registered to a reference image using FSL-FLIRT (normalized mutual information as cost function) to generate a motion corrected dynamic PET image series. The final PET image was computed as the average of the motion corrected images. The one minute frame length was required in order to have sufficient counts for the PET image reconstruction and image co-registration.

### PET image reconstruction

PET image reconstruction was performed with the following procedure. List-mode data were reconstructed with an ordinary Poisson ordered-subsets expectation maximization algorithm (OP-OSEM: 21 subsets, 3 iterations) and point spread function (PSF) correction using 344×344×127 matrix with a slice thickness 2.03 mm and a pixel size 2.09 mm. The reconstructed PET data were smoothed using a 3-D Gaussian filter (5 mm in all directions). The image reconstruction was implemented using the scanner software.

### Data acquisition

A total of eleven healthy human subjects were acquired on a MR-PET scanner (Siemens Biograph mMR, Erlangen, Germany) equipped with a 20-channel head and neck coil at Monash Biomedical Imaging, Melbourne, Australia. The human scans were approved by the Monash University human research ethics committee.

#### Motion controlled study

One healthy volunteer was injected with a bolus of 110 MBq [18-F]FDG, and instructed to move their head during the scan at specific times. Head motion was introduced during the EPI BOLD, as well as between structural scans (e.g. T1 and T2 weighted scans). MR images were acquired (see Table 2 for acquisition parameters). PET list-mode data were acquired for 60 minutes. The PET attenuation map was acquired using the UTE (ultrashort echo time) sequence on the Siemens Biograph mMR.

**Table 2:** Comparison of image registration bias (in mm) using T1-MPRAGE and T2-FLAIR images as references.

|  | **T1-MPRAGE** | **T2-FLAIR** |
| --- | --- | --- |
| **GRE-EPI** | $0.92 \pm 0.24$ | **$0.54 \pm 0.21$** |
| **Proton density** | $0.51 \pm 0.10$ | **$0.36 \pm 0.11$** |
| **SE-EPI** | $0.52 \pm 0.04$ | **$0.23 \pm 0.04$** |
| **2D GRE** | $0.96 \pm 0.17$ | **$0.39 \pm 0.10$** |

#### Group study – slow infusion based static and dynamic PET imaging

Ten subjects were administered 260 MBq FDG at a constant slow infusion rate of 36mL/hr over 90 minutes. MR images were acquired as per Table 2. Prior to data acquisition, the subjects were instructed to keep movements to minimum during the 90-min examination.

### PET image quality assessment

#### Static PET image reconstruction – Image sharpness

Relative image sharpness was used to quantify the motion correction improvements in the motion corrected PET images. The image sharpness was calculated using the mean absolute Laplacian of Gaussian (LOG) (Schleyer et al., 2015), defined as:

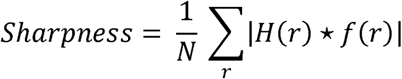

where *r* is the voxel index in the image *f(r)*, convolved with *H*, a 9×9×9 kernel describing the 3D LoG with standard deviation of 1.9. The group sharpness index was calculated as the mean and standard errors across the ten subjects in the slow infusion study.

#### Dynamic PET image reconstruction – DICE coefficients

To assess the quality of slow infusion dynamic PET data, DICE coefficients were calculated with the following steps. Firstly, the grey matter (GM) were segmented from the T2 weighted reference image using FAST with a probability threshold of 0.5 (Zhang et al., 2001). The DICE coefficients for the GM region were calculated for images reconstructed using the MR-guided MAF method, the fixed-MAF method, and the original motion corrupted images as follows:

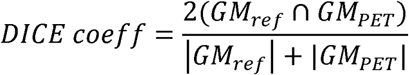

where *GM_ref_* is the grey matter mask from the T2 reference image, *GM_PET_* is the grey matter mask segmented from each of dynamic PET images [Hatt et al., 2009]. The operator ∩ is the intersection operator between two spatial masks. The DICE coefficient ranges from 0 (no spatial overlap between two images) to 1 (complete spatial overlap).

### Software availability

The MR-guided MAF method has been implemented in a fully automated software package, written in Python using the Arcana framework (Close et al. 2018), which is available as web-service: http://mbi-tools.erc.monash.edu/motion_correction. The source code for the package can be found at: https://github.com/MonashBI/banana/releases/tag/v0.2.0.

## Results

### Selection of MR reference image

The image registration imprecisions of the different MR image contrasts registered to the T1 weighted MPRAGE and the T2 weighted FLAIR, respectively, are compared in Table 2. The mean registration errors for both the T1 and T2 weighted images were less than 1mm. The optimal reference image was determined by comparing the mean registration errors for two contrasts, with theT2 FLAIR reference demonstrating a lower mean registration error based on the MR images acquired in this study.

### Motion controlled study results

The results for the single subject controlled head motion study, where the head movements were instructed multiple times during the examination, are given in Figures 2-4. The mean displacement plot (Figure 2) demonstrates a maximum movement close to 60 mm. Translation and rotation motion parameters are shown in supplementary Figures (S1a and S1b). The image from the MR-guided MAF with μ-map realignment shows symmetric radiotracer uptake in the two hemispheres of the brain (Figure 3a). Images reconstructed without μ-map realignment for the MR-guided MAF (Figure 3b), the fixed MAF (Figure 3c), and without motion correction (Figure 3d) demonstrate asymmetric radiotracer uptake in the brain hemispheres, which is also evident in line profiles drawn across the hemispheres in each image (Figure 3e). The asymmetric radiotracer uptake is almost certainly due to misalignment in sinogram space between the μ-map and the head position (see supplementary Figure S2). Using the sharpness index calculation, the images from the MR-guided MAF with μ-map realignment had a ∼23% increase in image sharpness compared to original motion corrupted images, and a ∼4% increase compared to the MR-guided MAF. Furthermore, the fixed-MAF images showed 21% greater blurriness compared with the fully corrected images.

**Figure 2.**
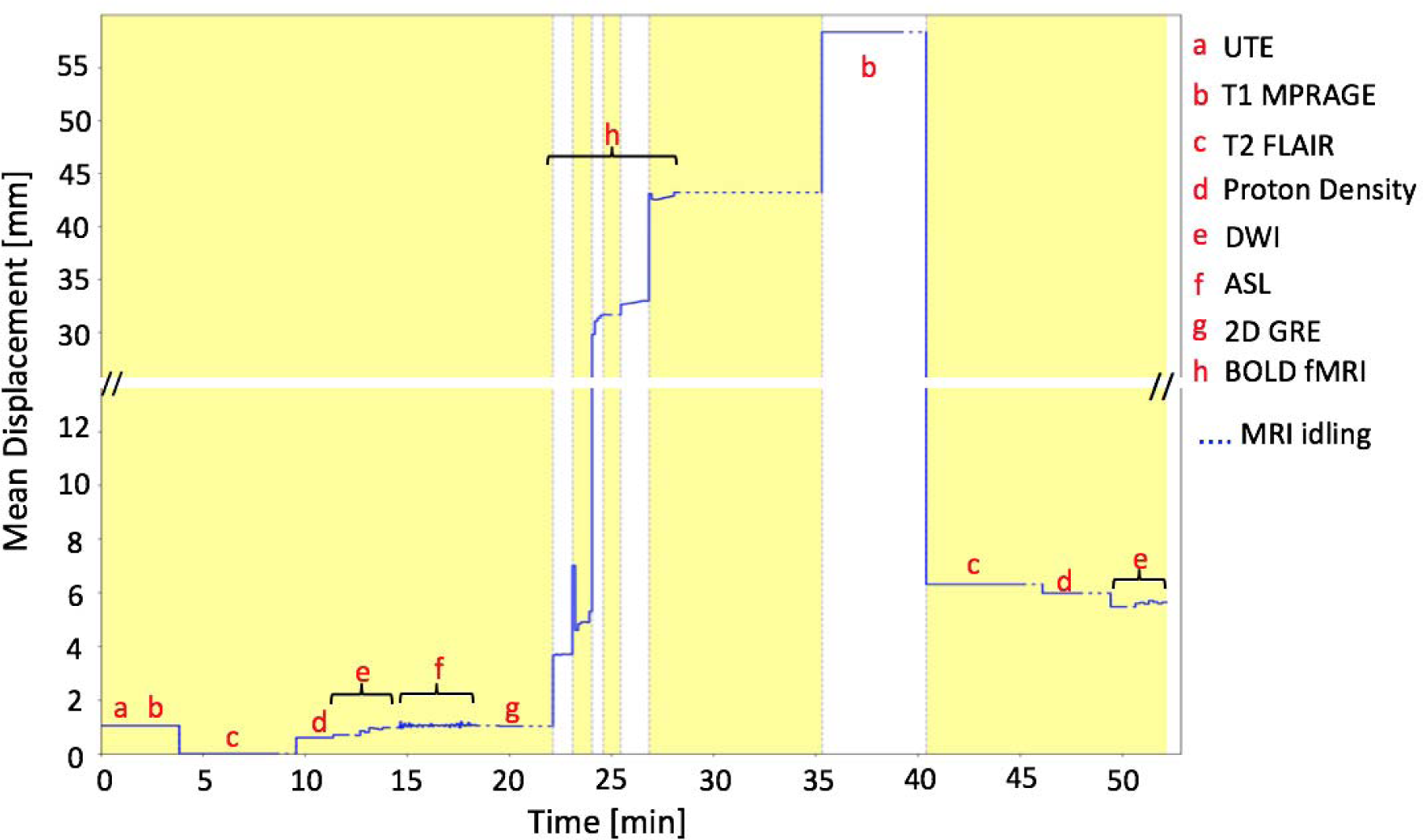
Mean displacement plot for the motion instructed volunteer demonstrating the head movement with respect to the T2 weighted reference image, as detected by the multi-contrast registration method. The yellow/white alternation bands indicate the durations of successive motion correction frames.

**Figure 3.**
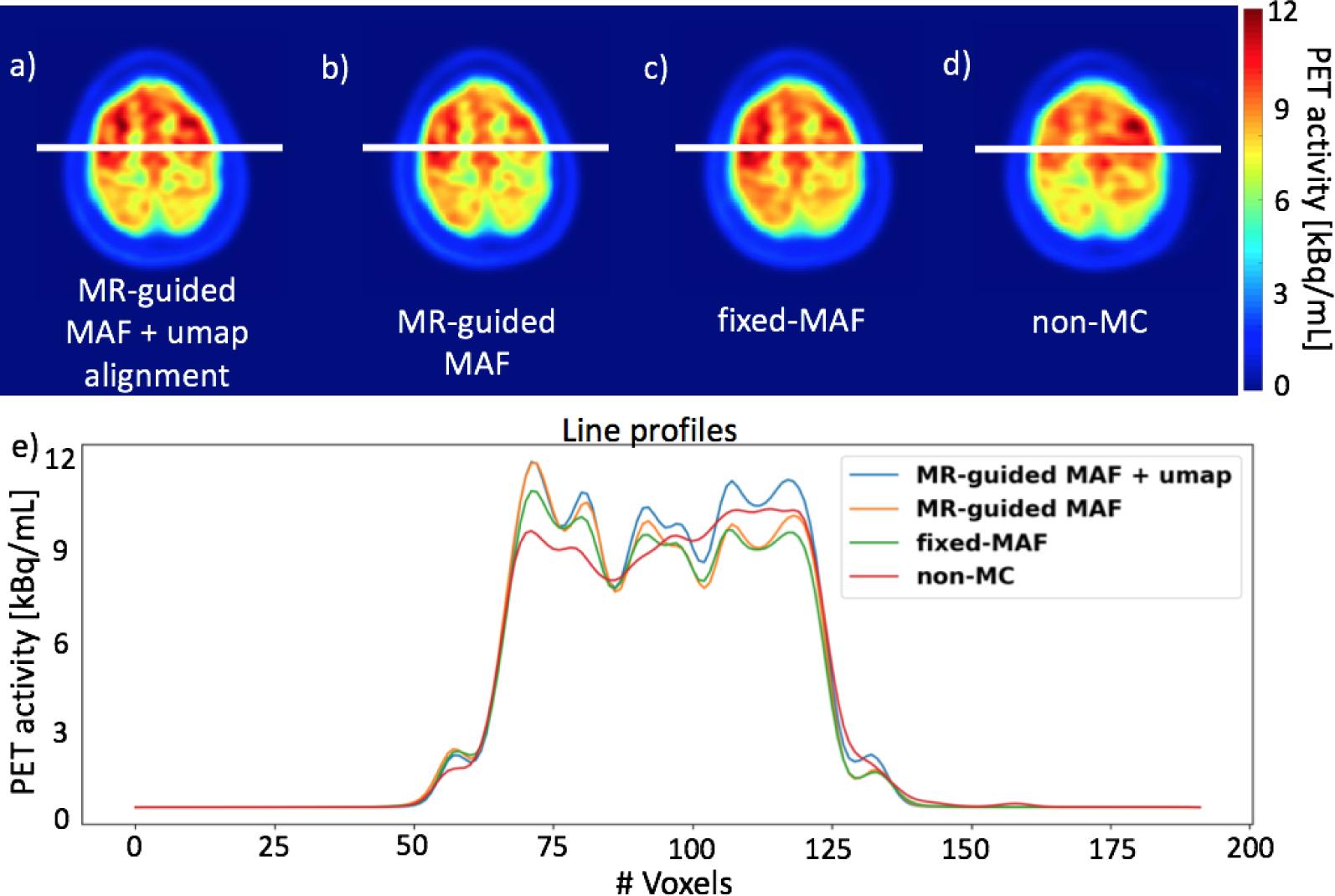
Motion correction results for the controlled motion experiment. Images in panels (a)-(d) show the reconstructed PET images using different reconstruction methods. The plots in panel (e) show the signal intensity variation along the line profiles in panel (a)-(d).

### Group study results

The group of ten participants were instructed to keep motion to a minimum during the 90-min long MR-PET scan. The estimated motion parameters are shown in Table 4. Overall, a 2-5 mm mean displacement was observed in the group. Figure 4 compares the group averaged results from different motion correction methods with the non-motion corrected images. For each method, the reconstructed images for all subjects were co-registered together to derive the group averaged image. The fully motion corrected image (i.e. MR-guided MAF with μ-map realignment, Figure 4a) depicts improved grey and white matter contrast compared with the fixed-MAF (shown in Figure 4b) and the non-motion corrected images (shown in Figure 4c). The comparison of line profiles (shown in Figure 4d) shows that the best grey and white matter delineation is observed from the fully corrected image. All three images show symmetric tracer uptake in both hemispheres, which is in agreement with the estimated relatively small mean displacements.

**Figure 4.**
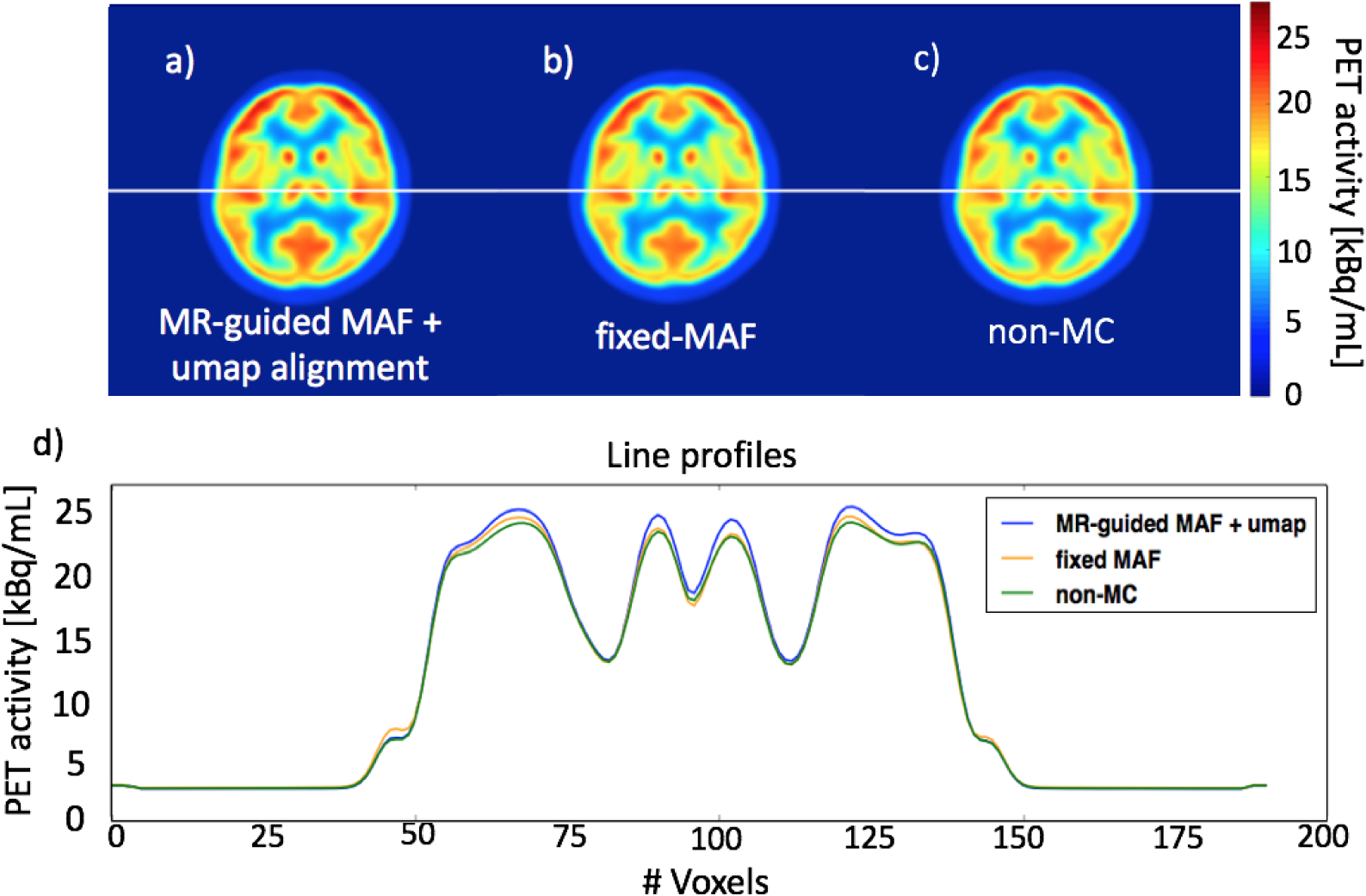
Comparison of the motion correction results for the group averaged image. The images in panels (a)-(c) show the reconstructed PET images using the three different reconstruction methods. The plots in panel (d) show the signal intensity variation along the line profiles in panels (a)-(c).

**Table 3:**
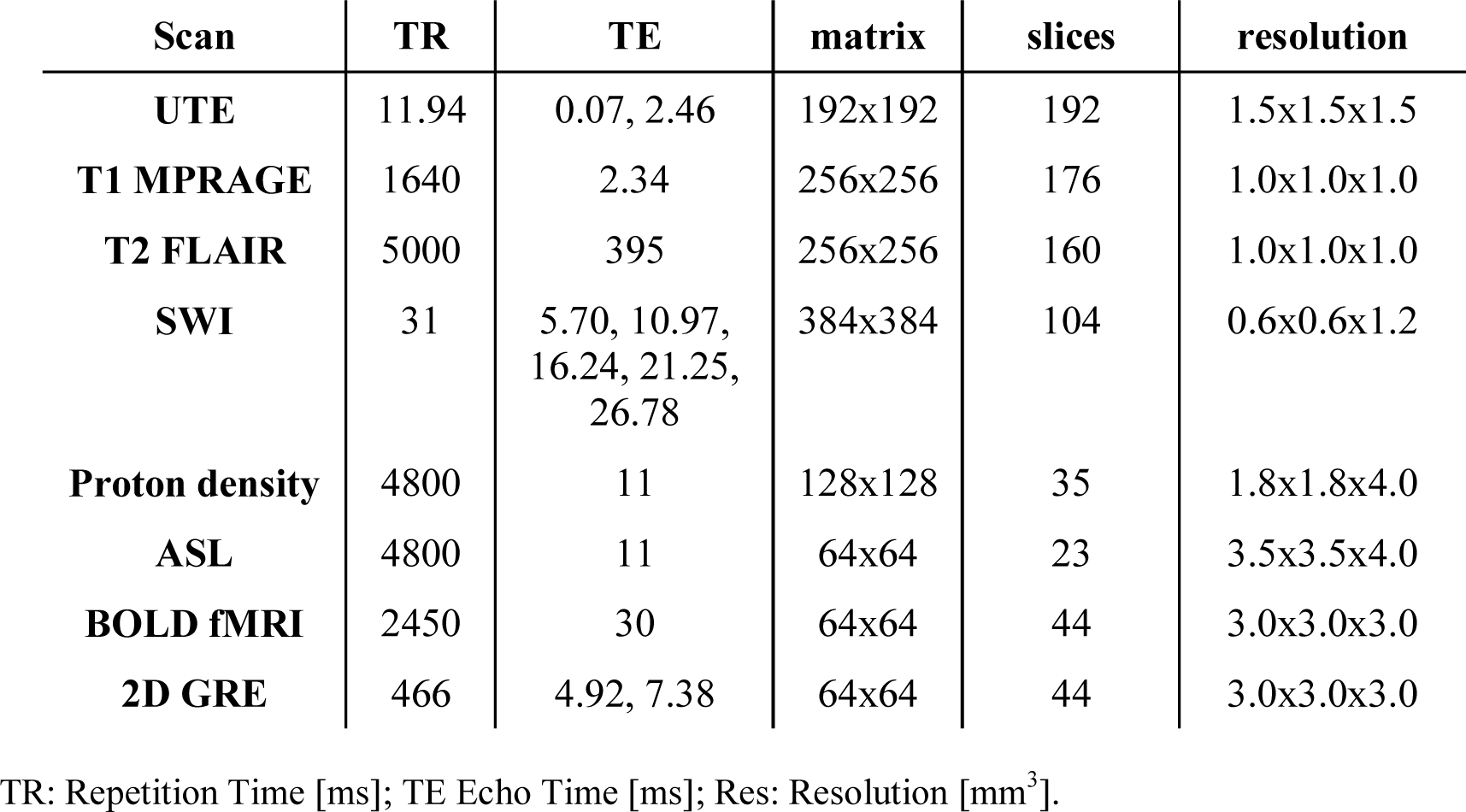
MR image acquisition parameters for the slow FDG infusion PET experiments.

**Table 4:**
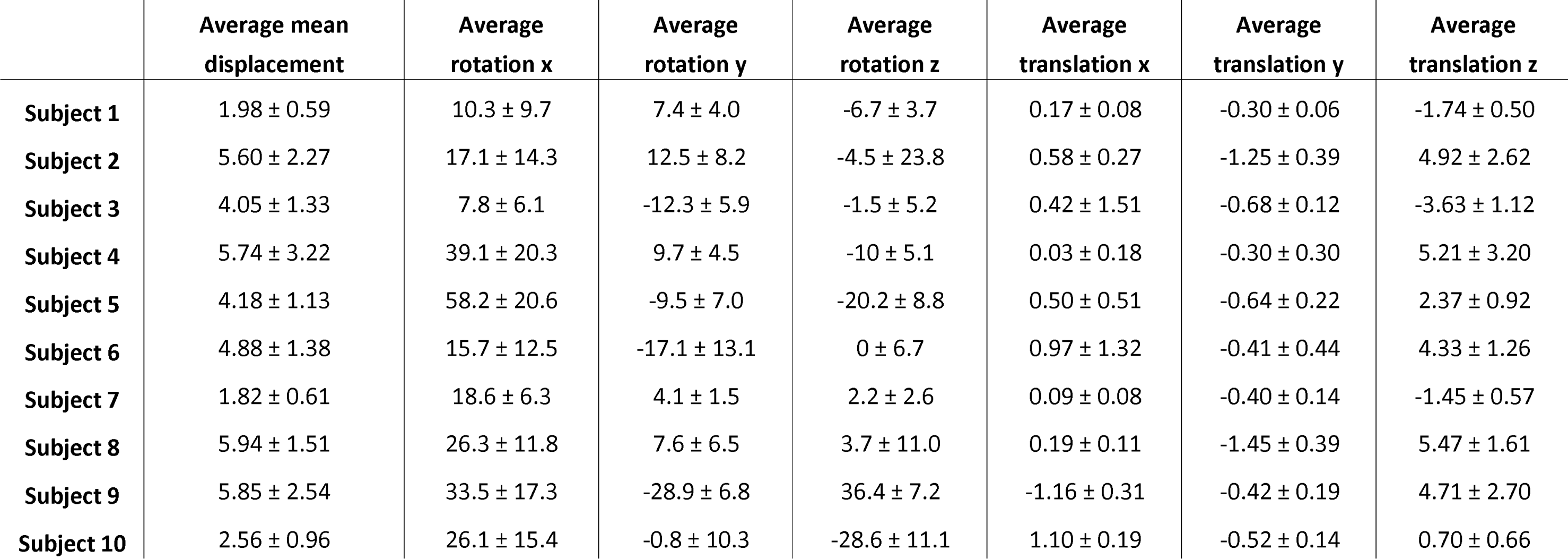
The average mean displacement [mm] and the average 6 motion parameters (3 for rotation [mrad] and 3 for translation [mm]) for the ten subjects used in group analysis.

|  | **Average mean displacement** | **Average rotation x** | **Average rotation y** | **Average rotation z** | **Average translation x** | **Average translation y** | **Average translation z** |
| --- | --- | --- | --- | --- | --- | --- | --- |
| **Subject 1** | 1.98 ± 0.59 | 10.3 ± 9.7 | 7.4 ± 4.0 | -6.7 ± 3.7 | 0.17 ± 0.08 | -0.30 ± 0.06 | -1.74 ± 0.50 |
| **Subject 2** | 5.60 ± 2.27 | 17.1 ± 14.3 | 12.5 ± 8.2 | -4.5 ± 23.8 | 0.58 ± 0.27 | -1.25 ± 0.39 | 4.92 ± 2.62 |
| **Subject 3** | 4.05 ± 1.33 | 7.8 ± 6.1 | -12.3 ± 5.9 | -1.5 ± 5.2 | 0.42 ± 1.51 | -0.68 ± 0.12 | -3.63 ± 1.12 |
| **Subject 4** | 5.74 ± 3.22 | 39.1 ± 20.3 | 9.7 ± 4.5 | -10 ± 5.1 | 0.03 ± 0.18 | -0.30 ± 0.30 | 5.21 ± 3.20 |
| **Subject 5** | 4.18 ± 1.13 | 58.2 ± 20.6 | -9.5 ± 7.0 | -20.2 ± 8.8 | 0.50 ± 0.51 | -0.64 ± 0.22 | 2.37 ± 0.92 |
| **Subject 6** | 4.88 ± 1.38 | 15.7 ± 12.5 | -17.1 ± 13.1 | 0 ± 6.7 | 0.97 ± 1.32 | -0.41 ± 0.44 | 4.33 ± 1.26 |
| **Subject 7** | 1.82 ± 0.61 | 18.6 ± 6.3 | 4.1 ± 1.5 | 2.2 ± 2.6 | 0.09 ± 0.08 | -0.40 ± 0.14 | -1.45 ± 0.57 |
| **Subject 8** | 5.94 ± 1.51 | 26.3 ± 11.8 | 7.6 ± 6.5 | 3.7 ± 11.0 | 0.19 ± 0.11 | -1.45 ± 0.39 | 5.47 ± 1.61 |
| **Subject 9** | 5.85 ± 2.54 | 33.5 ± 17.3 | -28.9 ± 6.8 | 36.4 ± 7.2 | -1.16 ± 0.31 | -0.42 ± 0.19 | 4.71 ± 2.70 |
| **Subject 10** | 2.56 ± 0.96 | 26.1 ± 15.4 | -0.8 ± 10.3 | -28.6 ± 11.1 | 1.10 ± 0.19 | -0.52 ± 0.14 | 0.70 ± 0.66 |

Figure 5 shows an averaged sharpness index (mean and standard errors) of the ten participants. The fully corrected images demonstrate a 7% increase in mean sharpness index when compared with fixed-MAF, and 12% increase when compared with non-motion corrected images. These differences are all statistically significant (***p<0.005, *p<0.05).

**Figure 5.**
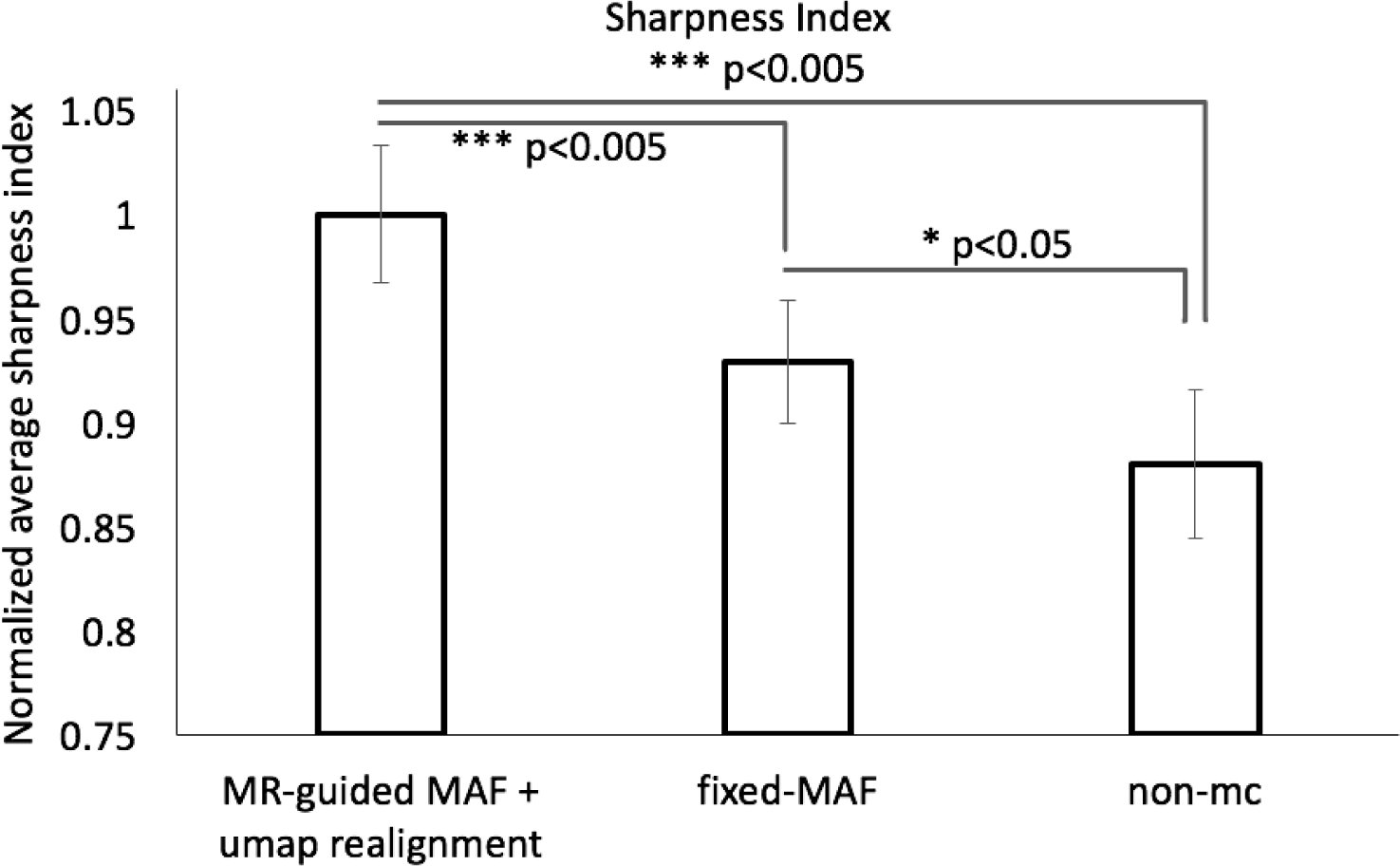
Comparison of the averaged (mean and standard errors) sharpness indices for the experimental group of ten participants.

### Dynamic PET image reconstruction

The segmented grey matter from a PET frame overlaid to the reference image was shown in Figure 6. The DICE coefficients were calculated and used to investigate the accuracy of motion correction in the grey matter region. Apart from the first 20 minutes, where the counts in the PET images were too low, the DICE coefficients of the MR-guided motion corrected images were constant around 0.65 (Figure 6b). The DICE coefficients of the frames aligned using the standard fixed-MAF approach, were lower compared with those calculated from the MR-guided MAF method. For both motion corrupted and fixed-MAF corrected images, the DICE coefficients fluctuated significantly toward the end of the 90-min acquisition where the head motion was greater (see mean displacement plot in supplementary Figure S3). In addition, the percentage difference in the time activity curve between the MR-guided MAF and the fixed-MAF varies between 1 to 5%, while that between the MR-guided MAF and the frames without motion correction reached 15% (see supplementary Figures S4a-b).

**Figure 6.**
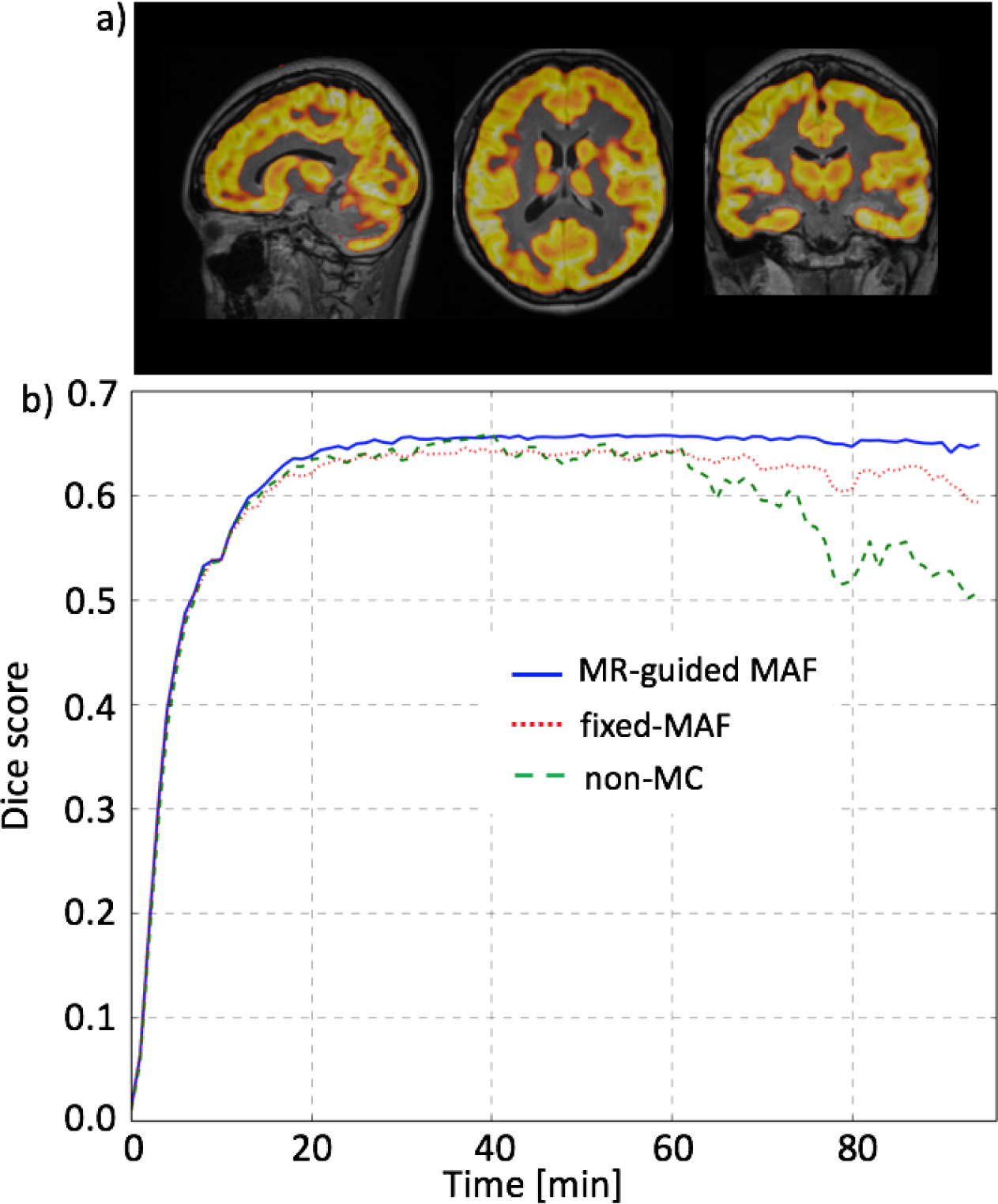
Comparison of the motion correction results between the MR-guided MAF, fixed-MAF and for the images without motion correction, for a dynamic PET reconstruction for one test subject. The Dice scores are shown in (b) using grey matter masks shown in (a).

The difference in the DICE coefficients between the non-motion corrected and the MR-guided motion corrected frames were significantly correlated with the mean displacement of the head position (Figure 7), demonstrating the accuracy of the MR-guided MAF approach.

**Figure 7.**
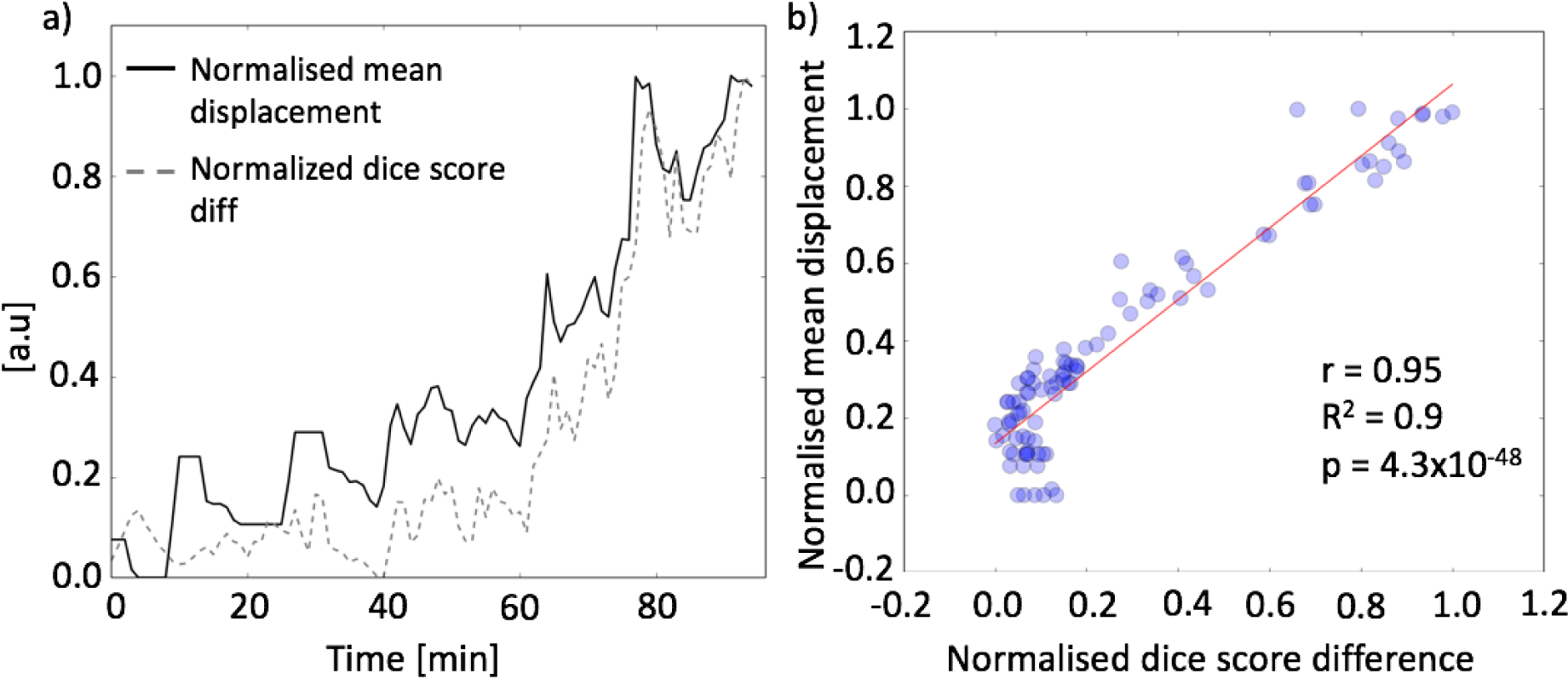
Plots of the Dice score differences and the mean displacement between the MR-based motion corrected and the non-motion corrected PET images in panel (a). Panel (b) shows the correlation scatter plot between the Dice score differences and the mean displacement.

## Discussion

The aim of this study was to develop a fully automated MR-based method to estimate and correct for head motion in simultaneous MR-PET imaging. The MR-guided MAF method relies on co-registration of multi-contrast MR images. One advantage of the MR-guided method is that no additional imaging navigators or dedicated EPI volumes are acquired for tracking motion (Keller et al., 2015) (Ullisch et al., 2012), thereby maintaining optimal usage of the scanning time, and simplification of the experimental workflow. Using both bolus injection and slow constant infusion FDG PET datasets, we have shown that the method removes head motion induced images artefacts, improved image sharpness, and provided more uniform tracer uptake across the brain. Compared with non-motion corrected images, the relative image sharpness increase using MR-guided MAF was ∼25% in the motion controlled study and an average of ∼12% in the subject cohort. The method using either MR-guided MAF with or without μ-map alignment performed better than the standard fixed-MAF, most likely because of intra-frame motion that is not corrected in the fixed-MAF method. The MR guided method can be expected to be robust and accurate even when the PET tracer activity is low, e.g. during the early phase of the slow infusion experiments and high temporal resolution dynamic PET image reconstruction (e.g. 20 secs).

Our findings highlight the importance of the re-alignment of the μ-map before PET image reconstruction if large motion occurs. Head motion causes a misalignment between the original head position during the attenuation map measurement and the head position during PET data acquisition. Consequently, without re-alignment of the μ-map, the reconstructed PET image has regions with inaccurate attenuation correction that lead to a significant quantification error of the tracer uptake. These inaccuracies can be recovered using the MR-aligned attenuation map, as shown by our results. In an [18-F]FDG PET dementia study, Chen et al., (2018a) used the time-weighted averaged coil μ-map to account for motion during image reconstruction. With the conventional fixed-MAF approach, re-alignment of the μ-map requires two reconstructions which can significantly increases the computation time and propagate the reconstruction errors.

The robustness and accuracy of the MR-guided MAF method has been demonstrated using datasets from a group of ten subjects. With motion between 2 to 5 mm, the MR-guided MAF achieves an increase in image sharpness around 7% compared with the fixed-MAF correction, and around 12% with respect to the non-motion corrected image. Compared with fixed-MAF, MR-guided MAF can more effectively correct intra-frame motion and requires fewer re-binned frames. Since the method forms fewer frames compared to the one minute binned fixed-MAF method, less computation time is required.

MR based motion correction has the advantage that the MR image co-registration is more accurate than using PET images due to the high spatial resolution and high contrast to noise ratio in MR images. Using both T1 and T2 weighted images as reference images, the image registration imprecision was less than 1mm, which is significantly less than the PET image resolution. Furthermore, PET data driven motion correction methods (Schleyer et al., 2015;

Thielemans et al., 2013) are heavily dependent on PET radioactivity count rates, which is problematic when dealing with short temporal frames and low dose applications. Conversely MR motion correction using EPI scans can provide a temporal resolution of two seconds or less. Several papers in the literature have used MR navigators to correct motion in PET imaging of the brain. EPI based fMRI were firstly used for motion navigators, and demonstrated improved PET image quality in several healthy subjects (Catana et al., 2011; Ullisch et al., 2012). In the same work, Catana et al., (2011) also implemented and evaluated cloverleaf MRI navigators (CLNs) to reduce motion artefacts. Instead of using dynamic EPI navigators, Keller et al (2015) inserted EPI volumes that were several minutes apart to exact motion information during the complete PET examination. Chen et al., (2018b) employed both fMRI navigators and EPI navigators that embedded inside the T1 weighted images to improve PET quantification accuracy in a group of FDG PET dementia patients. In our work, we further extended the MR based PET motion correction, and optimised for other MR contrasts (DWI/DTI, ASL and etc). This approach offers a motion correction strategy during the complete course of PET examination and for most popular MR neuroimaging sequences. The quantitative improvement in PET images was further evaluated in slow infusion based FDG PET datasets.

In our work, the motion correction has been implemented as a post reconstruction step (i.e. MAF), and there exist methods that apply motion correction prior or during image reconstruction. These methods are developed based on the consideration that better motion correction can be achieved at coincidence event level. Two early studies from Catana et al., (2011) and Ullisch et al., (2012) both applied motion correction to PET list-mode data prior to image reconstruction. They compared the list-mode motion correction with the post reconstruction based correction using the same motion estimates, and the final reconstructed images were found to be comparable. The advantage of applying motion correction prior to image reconstruction is that it can reduce the total number of reconstruction jobs, resulting in overall faster data processing. On the other hand, the advantage of using the MAF based method is the simple implementation and application in clinical and research MR-PET scanners. Although this work presents an MAF based motion correction, the motion parameters estimated using the multi-contrast MR image registration can potentially be fed into list-mode reconstruction. Jiao et al., (2016) proposed a method for joint estimation of PET kinetic parameters and correction of head motion during image reconstruction. Their method demonstrated improved accuracy in estimation of kinetic parameters, especially at low radioactivity doses and when large motion occurred, compared with the post reconstruction MAF method. Their work highlighted that PET data driven MAF methods suffer from inaccurate motion estimation when SNR is poor.

In this paper, we developed and applied the MR-guided MAF method to slow infusion dynamic PET imaging. Our results have shown the importance of motion correction for dynamic imaging approaches. Indeed, head movements can lead to large differences in the time activity curves, and the DICE score measures before and after the MR based motion correction. The difference between the time activity curves was correlated well with the motion estimates.

### Limitation and future work

The current method uses an MR derived attenuation map rather than a 68-Germanium based PET transmission scan or a CT X-ray derived attenuation map. Previous work by ourselves and others has demonstrated the accuracy of head image segmentation and the assignment of tissue attenuation values using advanced MR methods (Baran, et al., 2018). However, irrespective of the absolute accuracy of the attenuation maps used in the PET image reconstructions, the advantages of the fully motion corrected method (i.e. the MR-guided MAF with μ-map realignment) have been clearly demonstrated. One limitation of the current method is the absence of motion estimation during anatomical MR scans (e.g. T1 or T2 weighted). These scans can take several minutes, and motion may occur during these acquisitions. One possible solution is to insert navigator echoes (Tisdall et al., 2012) in the anatomical MR sequences. The navigators (image or k-space) can provide motion estimates in every hundreds of milliseconds to several seconds, depending on the repetition time of the sequence. Another possible solution is to use PET raw data driven motion correction during these MR sequences and during periods without MR data acquisition.

## Conclusions

In this paper, we have introduced a fully automated MR based motion correction method and software for simultaneous MR-PET imaging. Using *in vivo* datasets, the introduced MR-guided MAF method has shown significantly improved PET image contrast and sharpness compared with both non-motion corrected images and images from the conventional fixed-MAF method. The new method has also been applied to a slow FDG infusion dynamic PET study of brain metabolism, to produce significant improvements in PET image quality and accuracy of whole brain and regional time activity curve estimation.

## Acknowledgments

The research was supported by a grant from the Reignwood Cultural Foundation and an Australian Research Council (ARC) Linkage grant (LP170100494). GE is supported by the ARC Centre of Excellence for Integrative Brain Function (CE140100007). The authors acknowledge Richard McIntyre and Alexandra Carey for their assistance in acquiring data, Shenjun Zhong for assistance in software preparation, and Kamlesh Pawar for useful discussions. The authors acknowledge the useful discussions with Benjamin Schmitt, Thomas Gaass, and Daniel Staeb from Siemens Healthineers.

## Supplementary Figures

*Figure S1* Translational (a) and rotational (b) parameters for the motion instructed volunteer, as determined by the MR-guided MAF method.

*Figure S2*. Demonstration of the μ-map realignment in the MR-guided MAF method. Images in row (a) show misalignment between the head position and the original μ-map (highlighted in the circled area). The MR-guided MAF method re-aligned μ-map using the MR parameters completely recovers the PET image signal intensity as shown in row (b).

*Figure S3*. Mean displacement plot for a subject that shows the head movement with respect to the reference images, as detected by the MR-guided MAF method. The yellow/white alternation bands indicate the time durations for the motion correction frames.

*Figure S4*. Panel (a) shows the differences in the time activity curves as extracted from the MR-based motion correction method (blue), the fixed-MAF (red) and the non-motion corrected PET images (green). Panel (b) shows percentage errors between the whole head time activity curves extracted from the MR-based motion corrected frames and that from the fixed-MAF (grey line), and from the non-motion corrected (black line) PET images.

